# The gut microbiome and its metabolites are necessary for morphine reward

**DOI:** 10.1101/2020.09.17.302570

**Authors:** Rebecca S. Hofford, Nicholas L. Mervosh, Tanner J. Euston, Katherine R. Meckel, Amon T. Orr, Drew D. Kiraly

**Author notes:** Corresponding Author: Drew D. Kiraly, 1 Gustave L. Levy Pl, New York, NY 10029, USA.; Twitter: @kiralylab.

## Abstract

Recent evidence has demonstrated that the gut microbiome has marked effects on neuronal function and behavior. Disturbances to microbial populations within the gut have been linked to myriad models of neuropsychiatric disorders. However, the role of the microbiome in substance use disorders remains understudied. Here we show that animals with their gut microbiome depleted by non-absorbable antibiotics (Abx) exhibit decreased formation of morphine conditioned place preference and demonstrate marked changes in gene expression within the nucleus accumbens (NAc) in response to morphine. Replacement of short-chain fatty acid (SCFA) metabolites, which are reduced by microbiome knockdown, reversed the behavioral and transcriptional effects of microbiome depletion. This identifies SCFA as the crucial mediators of microbiome-brain communication responsible for the effects on morphine reward caused by microbiome knockdown. These studies add important new behavioral, molecular, and mechanistic insight to the role of gut-brain signaling in substance use disorders.

## Introduction

Opioid use disorder (OUD) is a devastating neuropsychiatric condition that leads to tremendous hardship for patients and families alike. In recent years overdose deaths from opioids have continued to rise, accounting for almost 70,000 lives lost last year in the United States alone^1,2^. The current state of the art treatment for OUD is the use of opioid agonist replacement therapies. While these therapies can be quite effective for some^3^, for too many patients they are ineffective or unpalatable. Additionally, they are associated with often intolerable side effects and carry the potential for dependence^4^ which can decrease compliance and contribute to relapse^5^. For this reason, pharmaceuticals and interventions focusing on non-traditional targets have gained interest as potential treatments or mitigation strategies for OUD.

Over the last decade, there has been increased interest in the resident population of bacteria in the intestinal tract, collectively called the gut microbiome, as a mediator of neuropsychiatric disease^6,7^. Individuals with autism spectrum disorder^8^, major depression^9^, and other neuropsychiatric conditions^10,11^ exhibit significantly disrupted gut microbial populations. While most clinical work is currently observational, preclinical studies have begun to clarify the mechanism of microbiome-brain crosstalk in neuropsychiatric disease. Based on this early work, the presence of a complete and diverse microbiome is necessary for normal brain function and behavior. Mice born germ-free (i.e. without any internal or external microbiome) exhibit alterations in anxiety-like and social behaviors^12,13^ as well as dysregulation of gene expression and chromatin structure in the brain^14,15^. Adult mice with their microbiome depleted with antibiotics exhibit changes in fear learning^16,17^, behavioral response to cocaine^18^, and corresponding transcriptional control in limbic brain regions^16–18^.

While there are multiple potential routes of gut-brain communication^19^, one of the best studied is via the production of neuroactive metabolites by the microbiome. One class of metabolite, the short-chain fatty acids (SCFA), which are formed by bacterial fermentation of dietary fiber^20^, have garnered particular interest as gut-brain signaling molecules. These molecules pass from the intestine into blood, can cross the blood-brain barrier^21^, and have been shown to affect development of neurodegeneration^22^, models of autism^23^, and behavioral response to cocaine^18^. SCFA serve a variety of cellular functions but have received attention for their ability to modify histone proteins and thus alter epigenetic regulation of gene expression. The three main SCFA, acetate, butyrate, and propionate, are known to act as histone deacetylase (HDAC) inhibitors^24^. Additionally, the SCFA can act as sources for histone acylations. Recently, acetate from the gut was shown to alter brain histone acetylation^25^, and butyrylation and propionylation are known histone modifications^26^. Regardless, by indirectly increasing histone acetylation, SCFA can influence gene expression^27,28^ and are a key potential signaling mechanism in the gut-brain pathway. Knockdown of the microbiome and reduction in SCFA has the potential to alter not only the epigenetic landscape but the transcriptional response to drugs of abuse, which in turn can modify behavior^29,30^.

Recently, our group demonstrated that alterations in the gut microbiome and SCFA influence drug reward for cocaine^18^. Reduction in the total number of bacteria and microbiome diversity through the addition of broad-spectrum antibiotics (Abx) into drinking H_2_O enhances cocaine conditioned place preference (CPP) at low doses^18^. Providing SCFA concurrent with Abx returns place preference to baseline levels, implicating the reduction in SCFA caused by microbiome depletion as responsible for the enhancement of cocaine CPP. Additional work demonstrated that the combination of opioids and gut microbiome disturbance can also affect cocaine reward. Morphine-dependent mice do not display cocaine CPP due to withdrawal-mediated anhedonia^31^. Interestingly, colonization of drug-naïve mice with the microbiome of a morphine-dependent mouse recapitulates the abolishment of cocaine CPP found in morphine-dependent mice^32^, suggesting that some aspects of opioid withdrawal might be attributed to the microbiome. However, it is currently unclear if microbiome depletion affects the rewarding properties of other drugs of abuse, such as opioids^7^.

While direct studies exploring the contribution of the gut microbiome to opioid reward or reinforcement have not been conducted, several pieces of evidence suggest interactions between the microbiome and some effects of opioids. Germ-free mice or mice with their microbiome knocked down via oral Abx do not develop tolerance to the antinociceptive effects of morphine^33,34^; this effect is reversed by colonization of the gut with a microbiome from healthy animals^33^. Additionally, Abx has been shown to disrupt the neuronal ensembles normally activated by oxycodone administration and withdrawal^35^, suggesting that the microbiome mediates opioids’ effects on both behavior and brain function.

The studies herein present the first direct test of the relationship between the gut microbiome and opioid reward using an oral antibiotic knockdown model. We find that mice lacking a complex microbiome have decreased development of preference for and sensitization to morphine. Additionally, we see that mice with a depleted microbiome have markedly different transcriptional and epigenetic responses in the nucleus accumbens (NAc) following opioid treatment. Finally, we see that repletion of SCFA metabolites to antibiotic treated animals reverses behavioral and transcriptional effects of microbiome depletion. Taken together, our findings provide evidence for a role of the microbiome in addiction-like behaviors and provide mechanistic insight into gut-brain signaling pathways.

## Results

### Oral Abx reduces microbiome diversity and alters the proportion of dominant bacterial populations in control and morphine-treated mice

Similar to our previous work^18^, we used non-absorbable broad spectrum antibiotics to induce a robust microbiome knockdown. This allows for interrogation of the effect of a deficient microbiome in a normally developed animal^36^. Mice were given 2 weeks of oral Abx delivered in their drinking water starting at 8 weeks of age before the start of once daily morphine injections (20 mg/kg, s.c.) for 7 days (**Fig. 1A**). This yielded four treatment groups: H_2_O-Sal, Abx-Sal, H_2_O-Mor, and Abx-Mor. As observed previously^18^, chronic Abx did not cause deleterious effects on animal health. Abx mice exhibited the same amount of weight gain (**Supplemental figure 1**). Abx also did not affect morphine metabolism or brain penetrance (**Supplemental figure 2**). However, as expected, Abx treatment significantly reduced alpha diversity as measured by the number of unique observed taxonomic units (OTUs, **Fig. 1B** – main effect of Abx: *F*_(1, 14)_ = 116.5, *p* < 0.0001; no effect of morphine: *F*_(1, 14)_ = 2.14, *p* = 0.17 and no interaction: *F*_(1, 14)_ = 0.14, *p* = 0.72) and by the Shannon diversity index (**Fig. 1C** – main effect of Abx: *F*_(1, 14)_ = 286.4, *p* < 0.0001; no effect of morphine: *F*_(1, 14)_ = 2.82, *p* = 0.12 and no interaction: *F*_(1, 14)_ = 0.01, *p* = 0.91).

**Figure 1.**
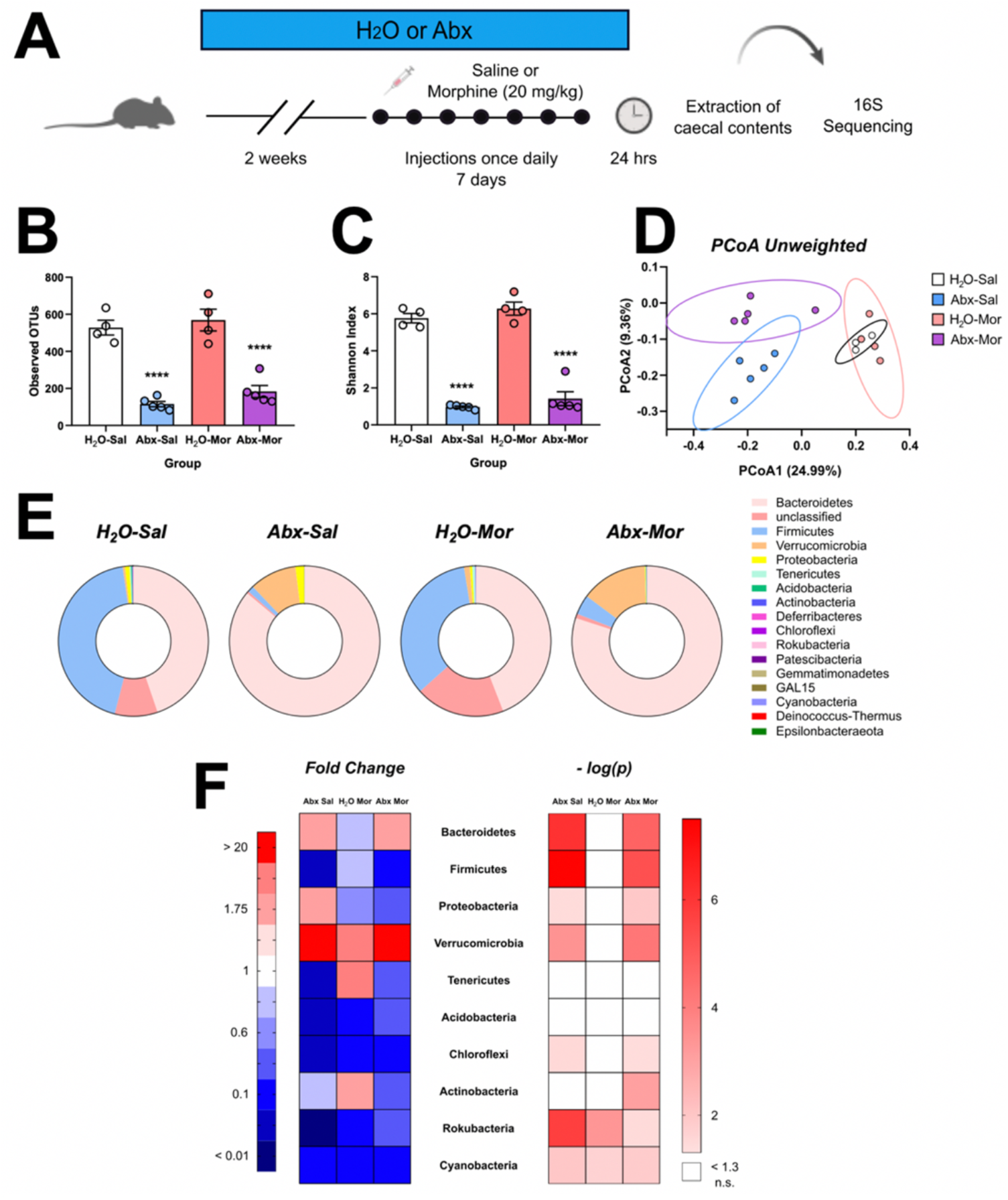
Oral Abx alters the microbiome. **(A)** Experimental timeline. **(B,C)** Abx reduces alpha diversity as measured by observed OTUs and the Shannon diversity index. **(D)** Mice from Abx-Sal and Abx-Mor possess unique microbiomes, but H_2_O-Sal and H_2_O-Mor populations overlap as measured using the unweighted Unifrac distance. **(E)** Relative phylum abundance in all groups of mice, each phylum shown in a different color. **(F)** Heatmap displaying the relative fold change (left) and -log(p) value (right) of the top ten most abundant phyla in control mice. Fold change values in red have greater abundance in that group compared to H_2_O-Sal and blue values have less abundance (left). Negative log(p) values in pink or red are significant (right). Data presented as means ± SEM. **** *p* < 0.0001.

Planned pairwise comparisons identified significant pairwise comparisons between H_2_O- Sal and Abx-Sal, H_2_O-Sal and Abx-Mor, and between H_2_O-Morphine and Abx-Mor (all *p* < 0.0001) for both measures.

However, morphine itself had minimal effects on microbiome composition, in contrast to other reports^32,33,37,38^. There was no difference between H_2_O- Sal and H_2_O-Mor on measures of alpha diversity (observed OTUs: *t*_(14)_ = 0.73, *p* = 0.47 and Shannon index: *t*_(14)_ = 1.21, *p* = 0.25). Beta diversity assessed using an unweighted UniFrac dissimilarity matrix shows near complete overlap between samples in H_2_O-Sal and H_2_O-Mor groups (**Fig. 1D**) and relative proportions of bacterial phyla were similar between H_2_O-Sal and H_2_O-Mor (**Fig. 1E**). Only two of the top ten most abundant phyla were significantly reduced by morphine alone: Rokubacteria (*t*_(7)_ = 8.08, *p* < 0.0001) and Cyanobacteria (*t*_(7)_ = 3.65, *p* = 0.0082); however, these phyla are expressed at very low abundance in all tested groups (both < 0.1%, **Fig. 1F**).

Abx, whether alone or in combination with morphine, also had a substantial effect on microbial composition as well as diversity. Both Abx-Sal and Abx-Mor groups had significant shifts in relative proportions of 7/10 major phyla, with the largest effects being a drastic reduction in Firmicutes (Abx-Sal: *t*_(7)_ = 26.12, *p* < 0.0001; Abx-Mor: *t*_(7)_ = 12.39, *p* < 0.0001) and expansion of both Bacteroidetes (Abx-Sal: *t*_(7)_ = 16.32, *p* < 0.0001, Abx-Mor: *t*_(7)_ = 10.27, *p* < 0.0001) and Verrucomicrobia (Abx-Sal: *t*_(7)_ = 6.39, *p* = 0.0004, Abx-Mor: *t*_(7)_ = 8.43, *p* < 0.0001), **Fig. 1F**.

Interestingly, the combination of Abx and morphine induced shifts in the microbiome that were different from both Abx-Sal and H_2_O-Mor. Abx-Sal and Abx-Mor are more similar to each other than the H_2_O groups based on phyla abundance (**Fig. 1E**), but sample clustering by Unifrac distance suggest these samples are from distinct populations (**Fig. 1D**). At the level of individual phyla, Actinobacteria was significantly changed in Abx-Mor only (Abx-Sal: *t*_(7)_ = 0.22, *p* = 0.83; Abx-Mor: *t*_(7)_ = 5.11, *p* = 0.0014) and Abx-Mor caused a reduction in the proportion of Proteobacteria (*t*_(7)_ = 3.40, *p* = 0.011) while Abx-Sal caused a significant increase (*t*_(7)_ = 2.55, *p* = 0.038, **Fig. 1F**). All fold change, *p* values, and mean % abundance are included in **Supplemental Table 1**.

### Bacterial enzymes that synthesize SCFA are predicted to be downregulated in Abx-Mor mice and SCFA levels are reduced in Abx-treated mice

Given the decrease in overall bacterial abundance found in Abx groups, we next focused on predicted functional enzymatic pathways that might be differentially affected by Abx and morphine. Phyla-level analysis indicated that there was a drastic reduction in the proportion of Firmicutes in Abx groups compared to control (**Fig. 1F**); many bacterial species in this phylum are producers of the SCFA butyrate^39,40^. We identified enzymes that are predicted to be involved in butyrate and propionate synthesis using Phylogenetic Investigation of Communities by Reconstruction of Unobserved States (PICRUSt2)^41^. Direct comparison of mean sequence proportion between Abx-Mor and H_2_O-Sal mice predicted multiple differentially regulated enzymes in Abx-Mor related to butyrate synthesis (**Fig. 2A**) and propionate synthesis (**Fig. 2B**)^39^. Full PICRUSt2 list is supplied as **Supplemental Table 2.**

**Figure 2.**
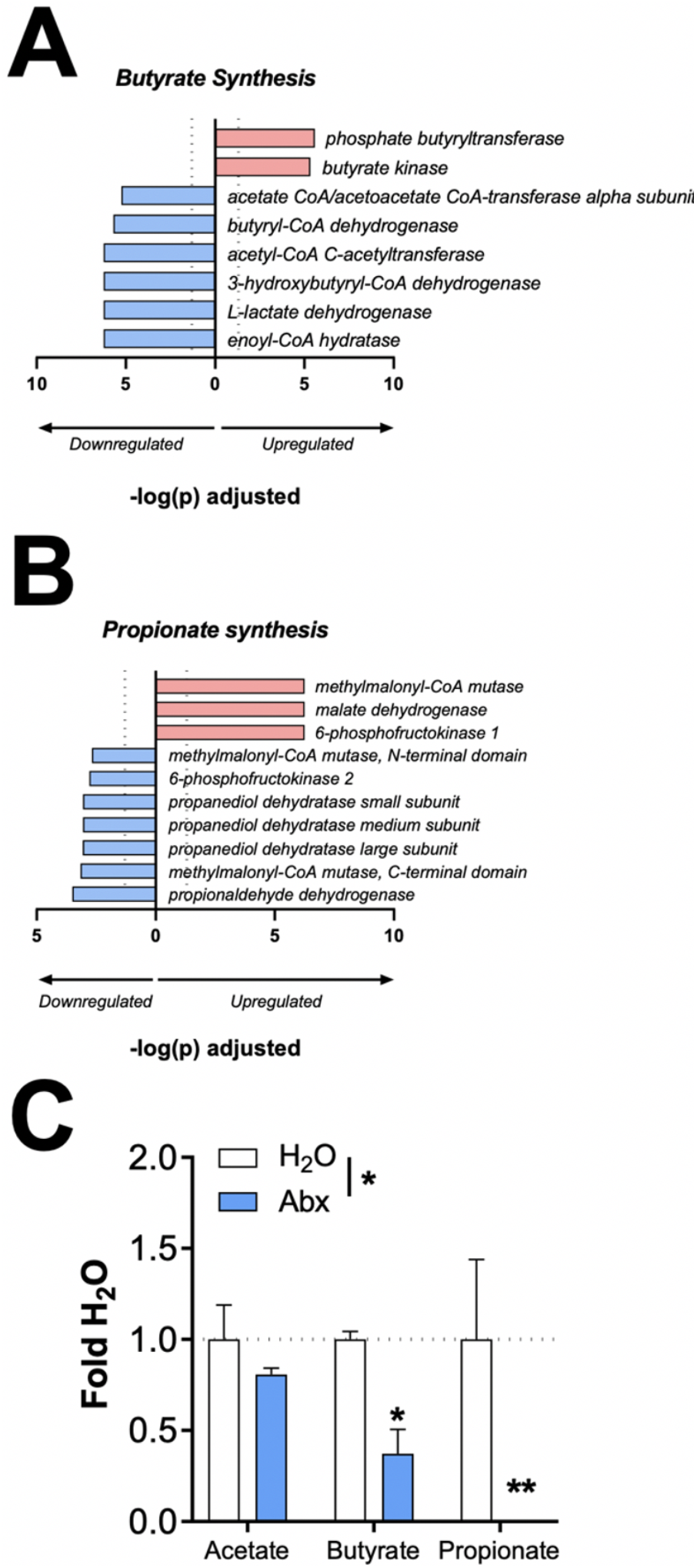
SCFA levels are reduced in Abx-Mor mice. **(A)** Enzymes that synthesize butyrate are mostly predicted to be downregulated in Abx-Mor mice. **(B)** Enzymes that synthesize propionate are predicted to be up- or downregulated in Abx-Mor mice. Pathways in pink are upregulated and pathways in blue are downregulated. Only significant results graphed. Horizontal dotted line at 1.3 indicates FDR-corrected significance level. **(C)** Levels of butyrate and propionate are significantly lower in the caecum of a separate group of Abx-treated mice. Data presented as means ± SEM.* *p* < 0.05, ** *p* < 0.01.

To confirm the predicted results from PICRUSt2, levels of the three most abundant SCFA - acetate, butyrate, and propionate - were quantified in the caecal contents of a separate group of Abx treated mice. Indeed, Abx significantly decreased levels of SCFA in caecal content (main effect of drink type : *F*_(1,8)_ = 29.41, *p* = 0.0006; no effect of metabolite: *F*_(2, 16)_ = 1.54, *p* = 0.24 and no interaction: *F*_(2, 16)_ = 1.54, *p* = 0.24). Planned comparisons between H_2_O and Abx mice found that butyrate (*t*_(24)_ = 2.18, *p* = 0.039) and propionate (*t*_(24)_ = 3.47, *p* = 0.002), but not acetate (*t*_(24)_ = 0.67, *p* = 0.51) were significantly reduced by Abx (**Fig. 2C**).

### Microbiome depletion reduces the persistence of morphine locomotor sensitization and morphine reward

To determine the behavioral effects of microbiome knockdown, mice were tested for morphine locomotor sensitization and morphine conditioned place preference (CPP). Both H_2_O and Abx animals developed locomotor sensitization to repeated morphine (main effect of day: *F*_(2.81, 22.50)_ = 7.83, *p* = 0.0011), but there was no effect of Abx (*F*_(1, 8)_ = 0.79, *p* = 0.40) and no interaction between day and Abx (*F*_(4, 32)_ = 1.45, *p* = 0.24), **Fig. 3A**. Upon morphine challenge given 10 days later, Abx mice had reduced persistence of locomotor sensitization compared to H_2_O mice (*t*_(8)_ = 2.73, *p* = 0.026, **Fig. 3B**).

**Figure 3.**
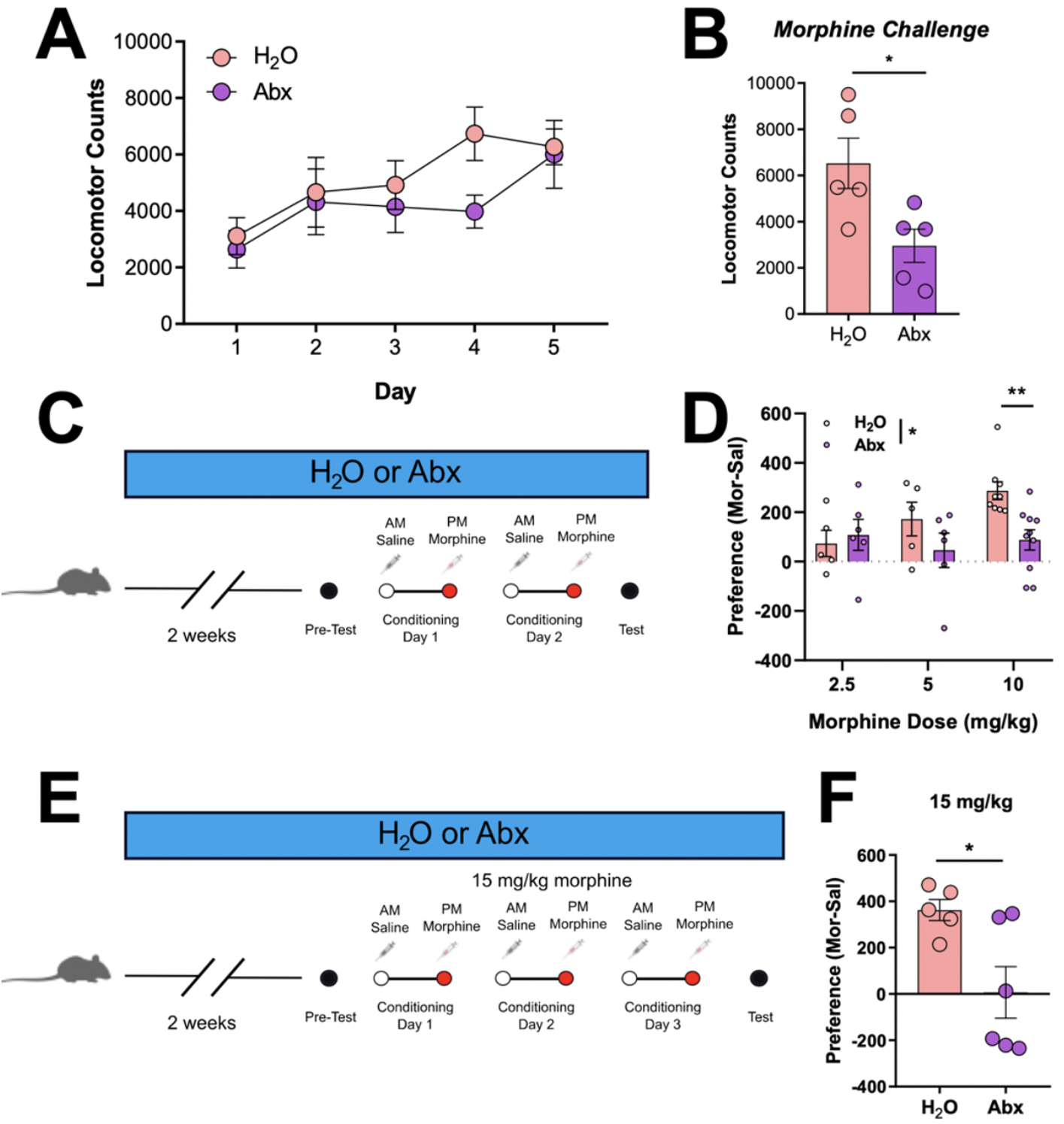
Microbiome knockdown reduces the persistence of locomotor sensitization and morphine place preference. **(A)** Abx did not alter the locomotor response to repeated morphine but did reduce the persistence of sensitization **(B)**. **(C)** Experimental timeline for 2-pairing CPP. **(D)** Abx treatment reduced high-dose morphine CPP. **(E)** Experimental timeline for 3-pairing CPP. **(F)** Abx mice did not demonstrate morphine CPP with 3 morphine-chamber pairings at a high dose (15 mg/kg). Data presented as means ± SEM. * *p* < 0.05.

Locomotor sensitization is often used as a tool for drug-induced neural plasticity, but it has little specificity as a model of reward. To better assess whether microbiome knockdown could influence opioid reward, mice from H_2_O and Abx cages were trained on morphine CPP. Conditioned place preference occurred over 4 days with 2 days of conditioning (**Fig. 3C**). On conditioning days, all mice were given injections of saline in the morning and 2.5, 5, or 10 mg/kg morphine in the afternoon. Microbiome knockdown reduced morphine place preference (main effect of drink type: *F*_(1, 35)_ = 4.76, *p* = 0.036; no effect of dose: *F*_(2, 35)_ = 2.13, *p* = 0.13 and no interaction: *F*_(2, 35)_ = 2.55, *p* = 0.09). Pairwise comparisons demonstrated that Abx mice specifically had reduced preference for the morphine-paired chamber at the 10 mg/kg dose (*t*_(35)_ = 3.17, *p* = 0.009), **Fig. 3D**.

To fully account for potential effects on dose response and changes in associative learning, we then performed enhanced morphine CPP conditioning with both an additional conditioning day (3 days vs 2 days) and a higher dose of morphine (15 mg/kg, **Fig. 3E**). Similar to their response at 10 mg/kg morphine, Abx mice had a reduced CPP score compared to H_2_O mice (*t*_(9)_ = 2.74; *p* = 0.023), **Fig. 3F**. Interestingly, the magnitude of effect *increased* as the dose of morphine increased (Cohen’s *d* 2.5 mg/kg = 0.26; 5 mg/kg = 0.78, 10 mg/kg = 1.04; 15 mg/kg = 1.73), suggesting that Abx more efficiently reduces the effects of high-dose opioids and this effect is unlikely due to impaired learning alone, since more morphine pairings potentiated the effect.

### Microbiome knockdown alters the NAc transcriptional response to morphine

While many brain areas contribute to drug reward and the development of addiction, the NAc is heavily implicated in primary reward. Lesions of this region abolish place preference and reduce ongoing self-administration^42,43^, making it an attractive neurobiological target to investigate molecular effects involved in the attenuation of morphine reward caused by microbiome knockdown. To explore the impact of microbiome depletion on gene expression in the NAc, H_2_O and Abx mice were given 7 daily injections of saline or morphine (20 mg/kg, s.c.) followed by NAc collection 24 hr after their last injection (**Fig. 4A**). RNA-sequencing revealed minor gene expression differences in mice receiving Abx alone compared to controls, with 51 transcripts differentially regulated in the Abx-Sal group (**Fig. 4B**). In contrast, there were 946 differentially regulated in the H_2_O-Mor group with the majority being increased in expression (**Fig. 4C**). However, the combination of Abx and morphine produced substantial gene expression differences that were greater than the effects of morphine and Abx alone, with 2806 genes downregulated and 2689 genes upregulated compared to controls (**Fig. 4D**). For full list of all significantly regulated genes in all comparisons, see **Supplemental Table 3.**

**Figure 4.**
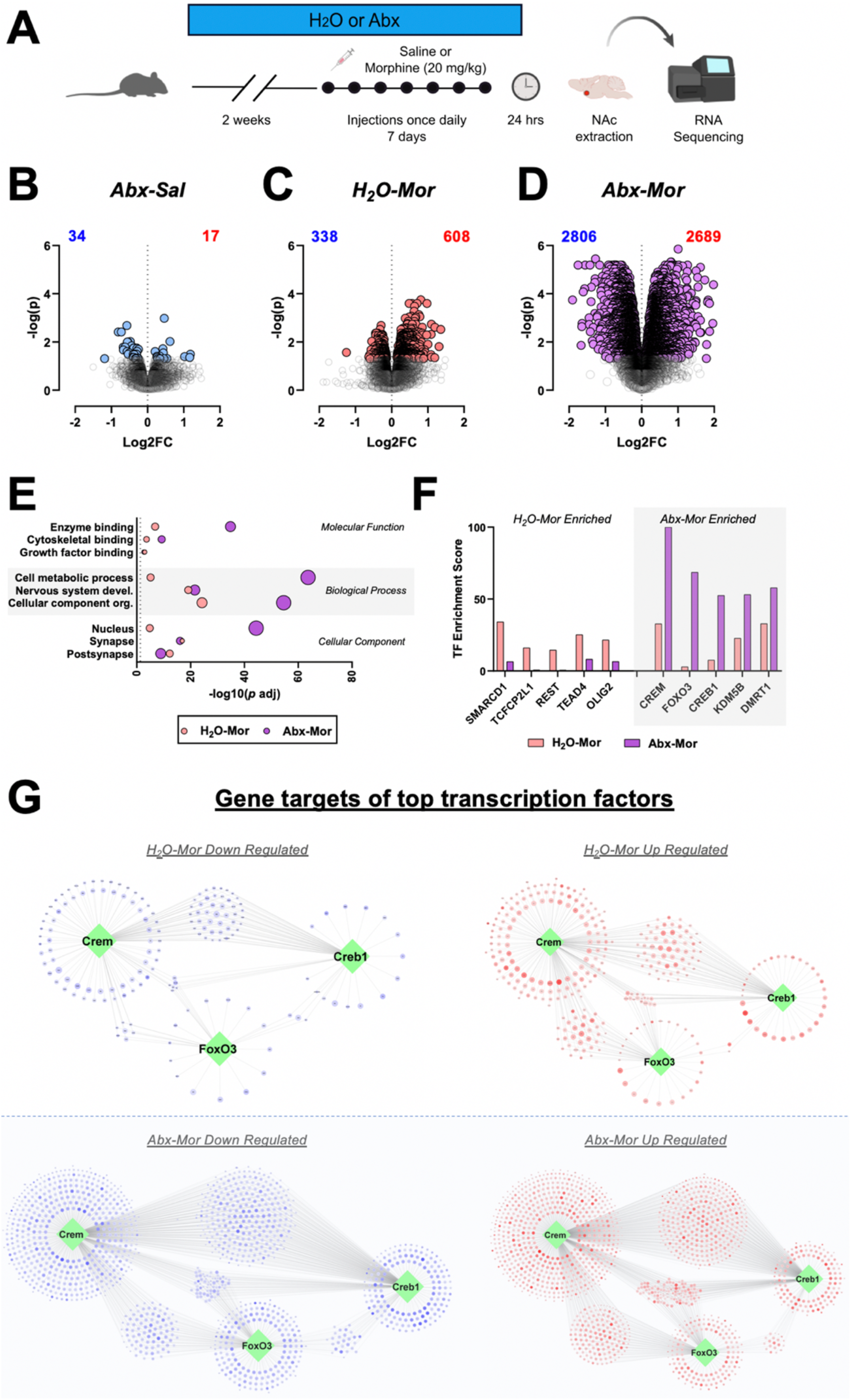
Morphine and Abx alter the NAc transcriptome. **(A)** Experimental timeline for RNA-sequencing. **(B-D)** Volcano plots depicting differential gene expression between H_2_O-Sal and Abx-Sal **(B)**, H_2_O-Mor **(C)**, and Abx-Mor **(D)**. Colored circles are significantly differentially regulated transcripts identified using an FDR corrected *p* < 0.05. **(E)** Representation of some top gene-ontology terms regulated in morphine groups. The x-axis is the FDR-corrected –log(*p*) value with the dotted line indicating an FDR corrected *p* level of < 0.05, and circle size represents number of genes per term. **(F)** Top differentially predicted transcription factors using Chea software analysis demonstrates factors with different predicted effects between groups (F). Y-axis is the overall transcription factor enrichment score predicted from genes in the dataset. Cloud diagrams depicting top regulated transcription factors (hubs) and their predicted downstream targets (nodes) (G).

To better understand how specific gene pathways were altered by morphine treatment in animals with and without a normal microbiome, we next performed Gene Ontology (GO) pathway analysis on significantly regulated genes from these three groups. We see that while morphine treatment regulated similar pathways in animals with and without a gut microbiome, the magnitude of the pathway effect was often greater in the Abx-Mor treated animals (**Fig. 4E**). Notably, Abx-Sal pathways are not depicted on the graph as none of these pathways had a -log(*p*) > 1.3 (i.e. not statistically significant) and thus would not show up on the graph. Abx-Mor animals had particular enrichment in pathways related to enzyme binding, cellular metabolism, and nuclear localization. However, H_2_O-Mor mice had more significant regulation of growth factor binding and post-synapse related genes, suggesting functional differences in how the two groups respond to the drug. Full pathway lists available for all comparisons in **Supplemental Table 4.**

Given the extent of the transcriptional changes between the two groups, we also sought to determine how changes in transcription factor activity might be playing a role in gene regulation in this model. Using data from the Chea transcription factor dataset, which utilizes ChIP-seq datasets to identify transcription factor binding peaks in the promoters of genes^44^ along with the Enrichr software package^45,46^, we identified transcription factors with the greatest differential transcription factor enrichment score between the H_2_O-Mor and Abx-Mor groups (**Fig. 4F**). We see that relative to genes in Abx-Mor mice, H_2_O-Mor mice had regulation of genes bound by Smarcd1, Tcfcp2l1 and Rest; whereas the top relative transcription factors predicted in Abx-Mor mice relative to H_2_O-Mor were Crem, Foxo3, and Creb1. Full data on transcription factor binding enrichment predicted via Enrichr are available as **Supplemental Table 5**. To better visualize the differences in predicted transcription factor activation patterns we performed analysis looking at up and down regulated genes downstream from the three transcription factors most enriched in the Abx-Mor group. As can be seen in **Fig. 4G** these transcription factors regulate networks of highly overlapping genes. However, in the H_2_O-Mor groups there are more distinct patterns of downstream genes that are primarily upregulated rather than downregulated. Whereas antibiotic treated mice show widespread up and downregulation of genes downstream from these factors – portending a much more widespread transcriptional dysregulation of these pathways in animals lacking a complex microbiome.

Given the differential behavioral response to morphine in animals with a depleted microbiome, we next investigated how microbiome depletion specifically affected the transcriptional response to morphine. When examining genes differentially regulated relative to H_2_O-Sal we see that there is considerable overlap between genes affected in the H_2_O-Mor and Abx-Mor groups (**Fig. 5A**), with approximately 77% of those genes regulated by morphine in the H_2_O group also being regulated in the Abx group. Further, when the direction of change was analyzed, we see that nearly all of these genes that were changed by morphine in both treatment groups were changed in the same direction (**Fig. 5B**). Gene ontology pathway analysis shows that genes similarly affected by morphine in both Abx and H_2_O groups had functionality related to protein binding, cellular organization, and synapse-related genes among others (**Fig. 5C**) full pathway analysis on overlapping genes available as **Supplemental Table 6**).

**Figure 5.**
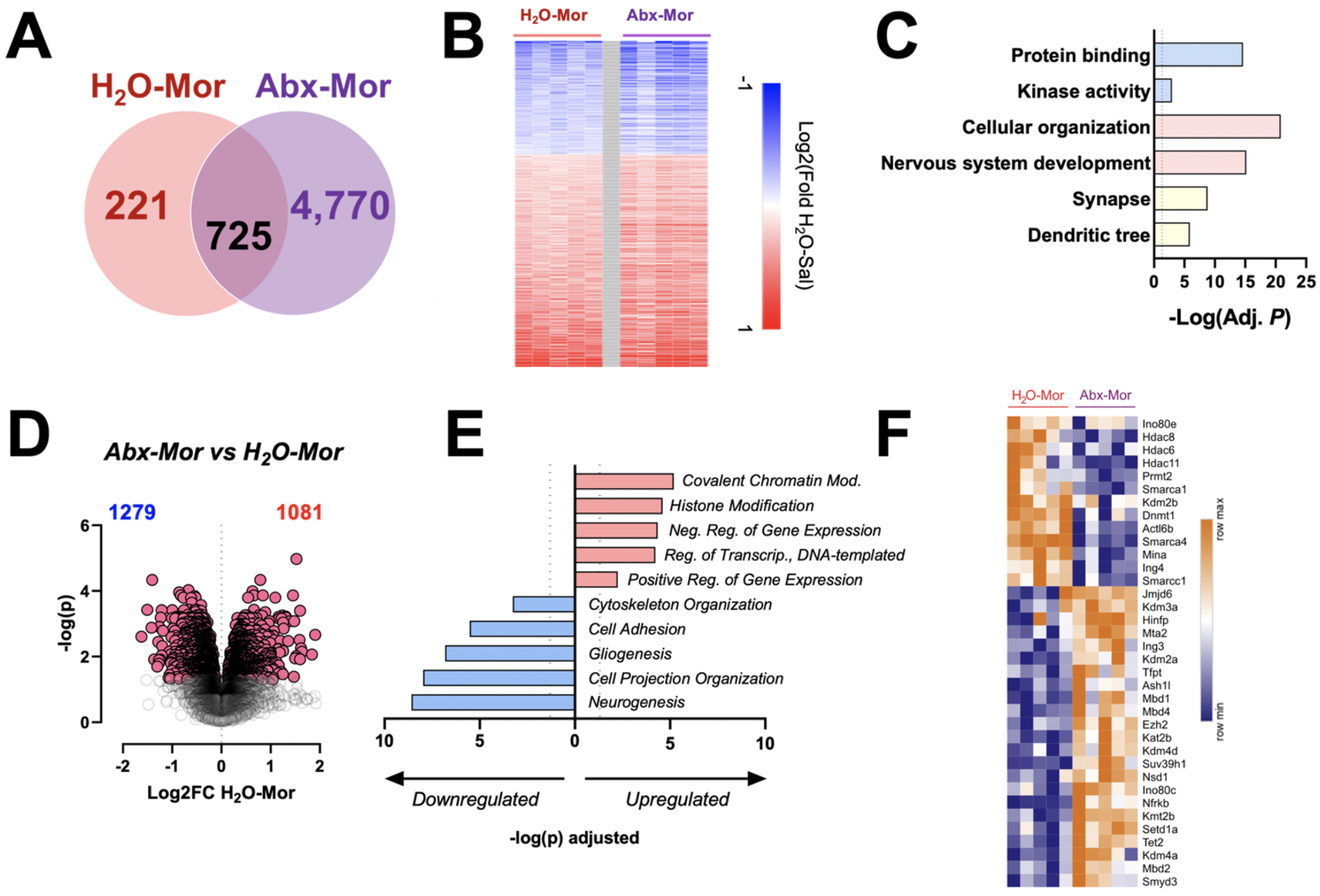
Microbiome knockdown alters the NAc transcriptional response to morphine. **(A)** Venn diagram depicting number of differentially regulated genes of H_2_O-Mor vs H_2_O-Sal (pink) and Abx-Mor vs H_2_O-Sal (purple). **(B)** Heatmap of genes that are differentially regulated by both H_2_O-Mor and Abx-Mor. **(C)** GO pathway analysis of overlapping genes. **(D)** Volcano plot depicting differential gene expression between Abx-Mor and H_2_O-Mor. Colored circles are significantly differentially regulated transcripts identified using an FDR *p* < 0.05. **(E)** Gene ontology pathways up and down regulated between Abx-Mor and H_2_O-Mor. Pink pathways are upregulated, and blue pathways are downregulated. Dotted vertical line is the significance cut-off of FDR *p* < 0.05. **(F)** Heatmap depicting all significantly regulated histone modifiers between Abx-Mor and H_2_O-Mor.

To further investigate how the microbiome alters responses to morphine, we performed transcriptomic analysis comparing the two morphine-treated groups. In direct comparison, Abx-Mor caused differential expression of 2360 genes (1279 downregulated, 1081 upregulated) when compared to H_2_O-Mor (**Fig. 5D**). Pathway analysis on genes differentially regulated between H_2_O-Mor and Abx-Mor identified several pathways related to epigenetic modification that were upregulated in the Abx-Mor group, including covalent chromatin modification, histone modification, negative regulation of gene expression, and positive regulation of gene expression (**Fig. 5E**). Downregulated pathways were more varied but included neurogenesis, cell projection organization, gliogenesis, and cytoskeleton organization (for full list of significantly regulated pathways, see **Supplemental Table 4**). To visualize the effect of Abx-Mor on the expression of histone modifiers, the Database of Epigenetic Modifiers (dBEM)^47^, a curated list of human epigenetic modifiers from healthy and cancerous genomes, was used to identify epigenetic regulators in the dataset (**Fig. 5F**), demonstrating that Abx differentially influences epigenetic modifier expression after morphine.

### Replenishment of SCFA reverses gene expression changes and morphine reward deficit caused by Abx

Since knockdown of the microbiome altered the behavioral (**Fig. 3**) and transcriptional responses (**Figs. 4 and 5**) to morphine in line with a reduction in short-chain fatty acid (SCFA) signaling (**Fig. 2**), we next examined if repletion of the SCFA via the drinking water could reverse the effects of microbiome depletion. For these purposes, animals received treatment with H_2_O, Abx, a cocktail of SCFA known to recapitulate physiological levels of SCFA^14,18,48^, or combination SCFA+Abx before examination of their effects on morphine CPP. Mice underwent the two-day conditioning protocol using 10 mg/kg morphine (**Fig. 6A**). Two-way ANOVA found no significant main effects (**Fig. 6B** - Abx: *F*_(1, 25)_ = 1.48, *p* = 0.23; SCFA: *F*_(1, 25)_ = 2.53, *p* = 0.12) or an interaction (*F*_(1, 25)_ = 3.33, *p* = 0.08). However, pairwise comparisons indicated that Abx reduced morphine place preference when compared to H_2_O as shown previously (*t*_(25)_ = 2.59, *p* = 0.016) and SCFA did not influence place preference by itself (SCFA vs H_2_O: *t*_(25)_ = 0.16, *p* = 0.87). However, SCFA supplemented mice with a reduced microbiome demonstrated normal morphine place preference (SCFA+Abx vs SCFA alone: *t*_(25)_ = 0.38, *p* = 0.71, **Fig. 6B**). This critical finding shows that the presence of SCFA metabolites can regulate behavior, even in the absence of a complex microbiome.

**Figure 6.**
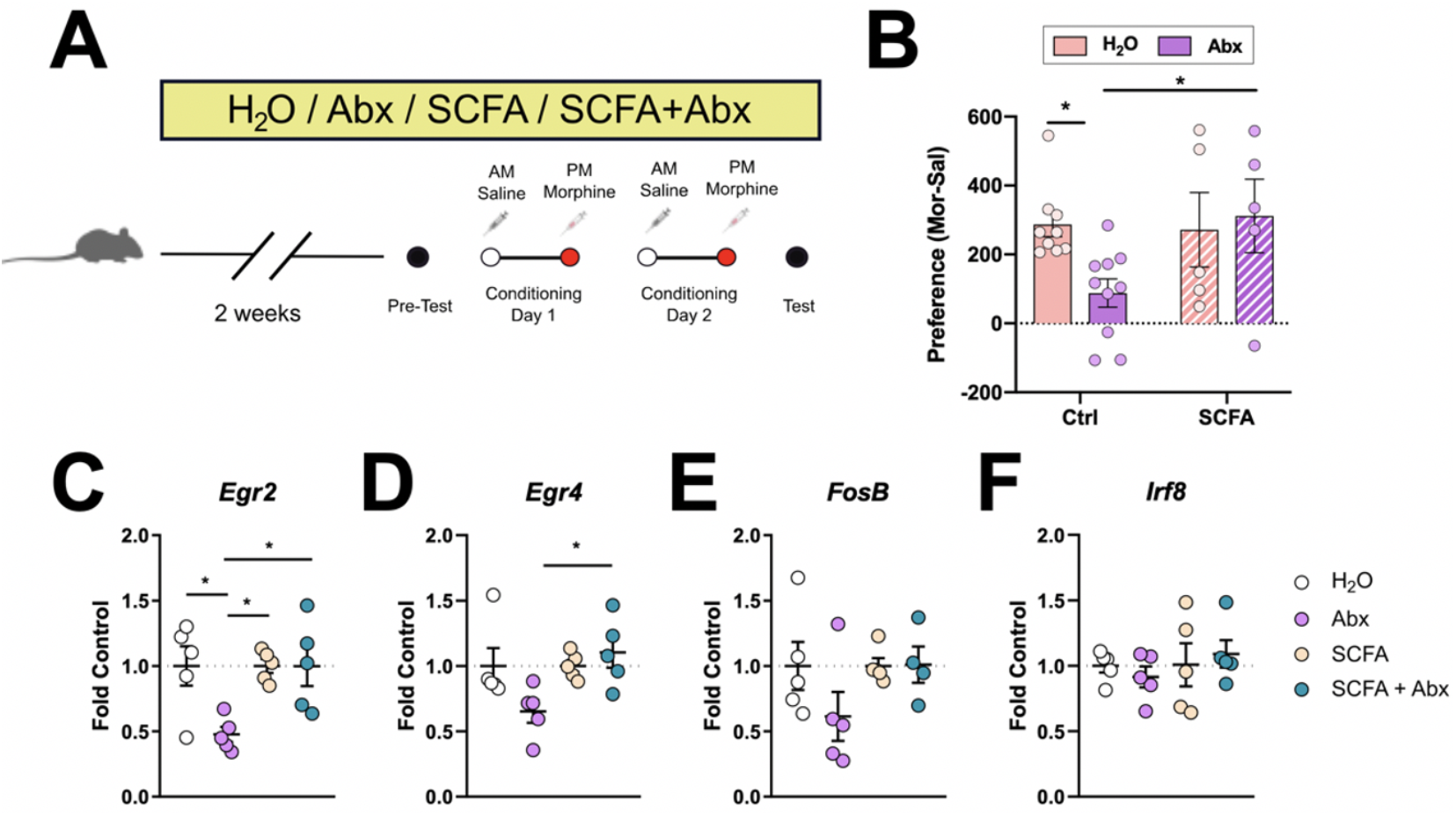
Replenishment of SCFAs reverses gene expression changes and morphine reward deficit caused by Abx. **(A)** Experimental timeline for CPP after SCFA replenishment. **(B)** SCFA restored morphine place preference in Abx-treated mice. SCFA+Abx normalized gene expression for *Egr2* **(C)** and *Egr4* **(D)** but did not change expression of *FosB* **(E)** or *Irf8* **(F)**. Data are presented as means ± Sem. * *p* < 0.05.

Using the same Abx and SCFA repletion paradigm, we then performed quantitative PCR analysis of gene expression levels in the NAc focused on targets known to be affected in the Abx-Mor groups from above. Here we found significant effects of drink type on multiple transcription factors including *Egr2* (main effect of Abx: *F*_(1, 16)_ = 5.27, *p* = 0.036, main effect of SCFA: *F*_(1, 16)_ = 5.23, *p* = 0.036, and a significant interaction *F*_(1, 16)_ = 5.23, *p* = 0.036, **Fig. 6C**) and *Egr4* (main effect of SCFA: *F*_(1, 16)_ = 4.87, *p* < 0.042 and a significant interaction: *F*_(1, 16)_ = 4.87, *p* < 0.42, **Fig. 6D;** no effect of Abx: *F*_(1, 16)_ = 1.43, *p* = 0.25), but no effect on transcript levels of *FosB* (Abx: *F*_(1, 16)_ = 0.80, *p* = 0.39; SCFA: *F*_(1, 16)_ = 2.64, *p* = 0.12; interaction: *F*_(1, 16)_ = 2.64, p = 0.12, **Fig. 6E**) or *Irf8* (Abx: *F*_(1, 16)_ = 0.0001, *p* = 0.992; SCFA: *F*_(1, 16)_ = 0.72, *p* = 0.41; interaction: *F*_(1, 16)_ = 0.61, p = 0.45, **Fig. 6F**). Importantly, SCFA+Abx affected transcription of both *Egr2* and *Egr4* differently than Abx alone; Tukey’s post-hoc analysis indicated that Abx alone reduced *Egr2* compared to H_2_O (*p* = 0.024), but there was no difference in *Egr2* expression between SCFA and SCFA+Abx (*p* > 0.9999, **Fig. 6C**). Additionally, Abx decreased expression of *Egr4* (non-significant, trending); but levels of *Egr4* were significantly increased in response to SCFA+Abx compared to Abx alone (*p* = 0.03, **Fig. 6D**). This indicates that, despite a reduction in microbial diversity via Abx, supplementation of SCFA back into animal’s drinking H_2_O can reverse expression patterns of several different transcription factors and suggests a mechanism for transcriptional effects seen in our full RNA-seq dataset.

## Discussion

The current set of studies demonstrate that a complete and complex microbiome produces metabolites (SCFA) that are necessary for morphine reward using a conditioned place preference model. Oral Abx successfully reduced the complexity of the gut microbiome (**Fig. 1B-C**) and reduced caecal SCFA content (**Fig. 2C**) but had no observed off-target health effects (**Supplemental Figures 1 and 2**). Using this model, we saw that knockdown of the microbiome attenuated morphine place preference (**Fig. 3D**) and the persistence of locomotor sensitization (**Fig. 3B**). While perturbations to the microbiome affect multiple physiological functions, our data suggest that the reduction in SCFA (**Fig. 2C**) due to microbiome knockdown drives the behavioral effects of Abx, since replenishment of SCFA to Abx-treated mice restored morphine CPP (**Fig. 6B**).

One well-documented role of SCFA is their ability to inhibit HDAC activity and to alter transcription factor activity^24,49–51^. Additionally, gut-derived short-chain fatty acids such as acetate can serve as a source for direct histone acylation in the brain^25,26^. Given the marked changes in behavioral and transcriptional response to morphine in microbiome-depleted animals and the ability of SCFA repletion to reverse these effects, we propose a model wherein a complex gut microbiome and its metabolites serve as a homeostatic regulator of experience dependent gene transcription in the brain. When the microbiome is depleted, robust external stimuli such as repeated exposure to drugs of abuse leads to dramatic changes in behavioral and transcriptional response. When gut metabolites such as the SCFA are returned, this homeostatic mechanism is restored and response to drugs of abuse or other external stimuli is restored.

These are certainly not the first findings to demonstrate that changes in the microbiome can affect transcriptional control in the brain. Interestingly, mice raised fully germ-free have altered expression at baseline of immediate early genes and other activity induced genes in the amygdala and prefrontal cortex, with more robust changes in gene expression occurring in response to an external stimulus^17,52^. Another recent paper that also utilized antibiotic depletion found robust changes in gene expression in response to reactivation of a fearful stimulus^16^. Interestingly, this same paper used single nucleus sequencing to demonstrate that knockdown of microbiome complexity leads to changes in gene expression in all cell types identified^16^. Other papers that have looked at sorted microglia have found that germ-free or antibiotic treated mice have robust changes to microglial gene expression and chromatin structure^14,15^. Future studies in our model of substance use disorder will seek to investigate cell-specific effects to tease apart the full mechanism underlying microbiome-morphine interactions.

CPP is a useful tool for assessing a drug’s rewarding value and requires an animal to associate a distinct environment with a drug’s effects^53^. Place preference can only occur if an animal successfully learns the environment - drug association and simultaneously finds that drug rewarding. In the current study, it is not clear whether the Abx-induced reduction in morphine CPP was due to a decrease in the subjective rewarding properties of morphine or an impairment in environment-drug association. In general, HDACs act to repress gene expression and tend to inhibit learning processes and HDAC inhibitors such as SCFA strengthen them^54^. Previous work has demonstrated enhanced morphine reward and sensitization after systemic sodium butyrate injections in microbiome-intact mice^55^ and a study showed potentiated heroin place preference after NAc HDAC inhibition^56^. These findings are in line with our current results.

Previous work on the microbiome and opioids has shown that depletion of the gut microbiome, or germ-free status, reduces the formation of opioid tolerance^33,34^. While our regimen of morphine does not produce any symptoms of physical dependence or withdrawal, when considering the translational implications of gut-brain interactions in opioid use disorder, these studies are an important parallel to ours. Given that morphine tolerance and opioid-induced hyperalgesia can be key drivers of pathological opioid use, the fact that the microbiome can alter both opioid tolerance and reward is a key component to understanding how gut-brain interactions can be targeted to reduce the burden of opioid use disorder in human subjects.

Interestingly, while our studies showed marked effects of microbiome depletion on brain and behavior, we saw no effects of morphine treatment on microbiome composition. This is interesting as several other studies have shown family and species level changes in microbiome composition following prolonged morphine^32,33,37,38^. While morphine has known effects on gastrointestinal motility and function, it is noteworthy that these previously published studies used alternate dose and administration methods than our current work. Based on this, it seems highly likely that opioid-induced changes in microbiome composition are dependent on interactions of timing, dose, and route of administration.

This paper demonstrates that the gut microbiome and its metabolites, the SCFA, are crucial mediators of morphine reward. Reduction of the microbiome attenuated morphine place preference, and this effect was reversed by repletion of SCFA. These studies also point to altered transcriptional control within the NAc as a potential mechanism linking the microbiome to morphine reward. Understanding how peripheral systems such as the gut microbiome interact with the brain to affect drug reward is an exciting avenue of research; these studies could aid in the development of safe mitigation strategies to be used in conjunction with maintenance therapy in the treatment of OUD.

## Methods

### Animals

Male C57BL/6 mice (7-9 weeks old, Jackson Laboratories) were group-housed (4-5 mice/cage) in a humidity and temperature-controlled colony room on a 24h light-dark cycle (lights on at 7:00). Drink solutions and food were available *ad libitum* throughout the entirety of all experiments. All animal procedures were approved by the Mount Sinai IACUC and all procedures conformed to the “Guide for the Care and Use of Laboratory Animals” (National Research Council 2010).

### Preparation and delivery of drink solutions

Cages were randomly assigned to H_2_O, Abx, SCFA, or SCFA+Abx conditions, depending on experiment. Abx cocktail contained 0.5 mg/ml vancomycin (Chem-Impex International #00315), 2 mg/ml neomycin (Fisher Scientific #BP266925), 0.5 mg/ml bacitracin (Research Products International #B3200025) and 1.2 μg/ml pimaricin (Infodine Chemical #7681-93-8) in H_2_O. SCFA solution contained 40 mM butyrate (#303410), 25.9 mM propionate (#P1880), and 67.5 mM acetate (#S5636) in H_2_O (all SCFA purchased from Sigma Aldrich); SCFA+Abx was a combination of both drink solutions. Mice from Abx, SCFA, and SCFA+Abx cages had their water replaced with their respective drink solutions 2 weeks before the start of experimental procedures and all mice remained on their drink solutions until the conclusion of the study. Mice were weighed before placement on drink solutions and no less than once weekly thereafter.

### Morphine

Morphine sulfate was provided by the NIDA drug supply program from National Institute on Drug Abuse and was diluted in saline and injected at a volume of 10 ml/kg.

### Detection of morphine in serum and brain

Mice from H_2_O or Abx cages were given a single injection of 20 mg/kg morphine or saline (s.c.) and were rapidly decapitated 30 minutes later for collection of blood and brain. Dorsal striatum (DStr) was identified as the area immediately ventral to the corpus callosum on slices containing the anterior commissure and was removed using a 14 gauge blunt needle. DStr punches were flash frozen on dry ice and stored at −80° C. Blood was allowed to clot at room temperature for 1 hr before centrifugation at 1500x*g* for 15 mins at 4° C. Supernatant was collected and stored at −80° C. DStr punches were homogenized on ice in lysis buffer (100 mM Tris-HCl, 2 mM EDTA, 125 mM NaCl, and 1% Triton-X in H_2_O) before centrifugation at 25000x*g* for 15 mins. Resulting supernatant was used to assess amount of total protein using Pierce BCA Protein Assay Kit (ThermoScientific) per assay instructions.

A morphine ELISA kit (Abnova #KA0935) was used to quantify morphine in serum and DStr. All samples were run in duplicate; blood serum was diluted 1:50 in PBS before loading and 16 μg (1 μg/μl) of protein was loaded for each DStr sample well. Assay was run per given instructions and was the plate was read at 450nm and 650nm 30 mins after addition of stop solution.

### 16S sequencing

Mice from H_2_O or Abx cages were given once daily injections of 20 mg/kg morphine or saline (s.c.) for 7 days and were rapidly decapitated 24 h after their last injection. Caecal contents were removed, flash frozen on dry ice, and stored at −80° C. DNA from caecal contents were isolated using DNeasy PowerSoil Kit (Qiagen) following kit instructions with an extended bead beating step. DNA concentration was determined with a NanoDrop1000. A concentration of 100 ng/μl DNA was sent to LC Sciences (Houston, TX) for sequencing. PCR amplification was achieved using primers (341F/805R) targeting the V3 and V4 region of the 16S rRNA region of the bacterial genome and was sequenced on an Illumina NovaSeq (2 x 250 bp paired-end). Amplicons were chimera filtered, dereplicated, and paired-ends were merged using Divisive Amplicon Denoising Algorithm 2 (DADA2)^57^, which identified unique genomic features. These features were used to determine observed taxonomic units (OTU), defined as sequences with ≥ 97% similarity. OTU counts were used to determine the Shannon index of alpha diversity and principle coordinates analysis plots were generated using the Unifrac distance as an assessment of beta diversity using QIIME2 software^58^. OTUs were identified by comparing their genetic sequences to reference bacterial genomes using SILVA (Release 132)^59^ with confidence set to 0.7. The predicted functional profiles of microbial taxa were assessed using the Phylogenetic Investigation of Communities by Reconstruction of Unobserved States (PICRUSt2) software package^41^ using default settings. Sequence counts of all KEGG orthology (KO) pathways for each sample were converted to mean proportion percent using the formula: (sequence count for specific KO pathway / total sequence count of all pathways x 100). Mean proportion percent of KO pathways involved in SCFA metabolism were pre-selected and compared via FDR corrected t-tests.

### Detection of SCFA levels

To quantify caecal SCFA mice were placed on Abx or kept on H_2_O for 2 weeks before rapid decapitation and removal of caecal contents as described above (n = 4 – 5 / group). Short-chain fatty acids were quantified using a Water Acquity uPLC System with a Photodiode Array Detector. Samples were analyzed on a HSS T3 1.8 μm 2.1×150 mm column with a flow rate of 0.25 mL/min. The volume of injection was5 uL, the column held at 40°C, of the sample at 4°C, for a run time of 25 minutes per sample. Eluent A was 100 mM sodium phosphate monobasic, pH 2.5; eluent B was methanol; the weak needle wash was 0.1% formic acid in water; the strong needle wash was 0.1% formic acid in acetonitrile, and the seal wash was 10% acetonitrile in water. For Eluent A the gradient was 100% eluent A for 5 min, gradient to 70% eluent B from 5-22 min, and then 100% eluent A for 3 min. The photodiode array was set to read absorbance at 215 nm with 4.8 nm resolution. Samples were quantified against standard curves of at least five points run in triplicate. Standard curves were run at the beginning and end of each metabolomics run. Quality control checks (blanks and standards) were run every eight samples. Concentrations in the samples were calculated as the measured concentration minus the internal standard; the range of detection was at least 1 – 100 μmol/g stool.

### Locomotor Sensitization

Mice from H_2_O or Abx cages (n = 5 / group) were assessed for their locomotor response to repeated morphine using San Diego Instruments Photobeam Activity System. The locomotor arena consisted of a frame crossed with infrared beams in the *x* and *y* dimensions. Clean, empty rat cages were placed within this frame to contain the mice while simultaneously allowing penetration of the infrared beams. Animals could freely move throughout the space and infrared beam breaks were counted to determine locomotor activity. Abx and H_2_O mice were injected with 10 mg/kg morphine (s.c.) and were allowed to ambulate for 45 mins. This occurred once daily for 5 days. Ten days after the last morphine injection, mice were given a morphine challenge injection and were placed back into the locomotor arena to measure persistence of morphine locomotor sensitization.

### Morphine conditioned place preference

Mice from H_2_O, Abx, SCFA, or SCFA+Abx (n = 5 – 10 / group) underwent morphine conditioned place preference (CPP) largely as described previously^18^ using Med Associates automated boxes and software. Each CPP apparatus had 3 distinct chambers with one small entry chamber and two larger conditioning chambers on either side of the entry room. End chambers were distinct in wall color and floor texture. The left end chamber had gray walls and a large grid floor, and the right end chamber had black and white striped walls and a small grid floor. Conditioned place preference occurred over 4 or 5 days; day 1 was pre-test, days 2-3 (or 2-4) were conditioning days, and the last day was test. On pre-test day, mice were placed into the center chamber of the apparatus and were allowed to explore all 3 chambers for 20 mins. Time spent in each chamber was recorded; mice spending > 70 % of their time in one chamber were excluded. Mice were assigned to their morphine-paired chamber using an unbiased approach such that half of each group had morphine injection paired with gray chamber. On conditioning days, mice were injected with saline (s.c.) in the morning and confined to one end chamber and injected with either 2.5, 5, 10, or 15 mg/kg morphine (s.c.) and confined in the opposite end chamber in the afternoon. Both morning and afternoon conditioning sessions lasted 45 mins. On test day, mice were again allowed to explore all 3 chambers of the apparatus for 20 mins, as described above for pre-test. Place preference score was calculated as: (time spent in the morphine-paired chamber on test day) – (time spent in saline paired chamber on test day).

### RNA-sequencing of NAc

H_2_O and Abx mice (same mice used for 16S sequencing, n = 5 / group) were given once daily injections of 20 mg/kg morphine or saline (s.c.) for 7 days and were rapidly decapitated 24 h after their last injection. Nucleus accumbens was identified by the anterior commissure and removed using a 14 gauge blunt needle. NAc punches were flash frozen on dry ice and stored at −80° C. RNA was extracted from the tissue using Trizol according to standard procedures. The integrity and purity of total RNA were assessed using Agilent Bioanalyzer and OD260/280 using Nanodrop. cDNA was generated using Clontech SMARTer cDNA kit (Clontech Laboratories, Inc., Mountain View, CA USA, catalog# 634925) from total RNA. cDNA was fragmented using Bioruptor (Diagenode, Inc., Denville, NJ USA), profiled using Agilent Bioanalyzer, and subjected to Illumina library preparation using SPRIworks HT Reagent Kit (Beckman Coulter, Inc. Indianapolis, IN USA, catalog# B06938). The quality and quantity and the size distribution of the Illumina libraries were determined using an Agilent Bioanalyzer 2100. The samples were then sequenced on an Illumina HiSeq2500 which generated paired-end reads of 106 nucleotides (nt). Data generated and checked for data quality using FASTQC (Babraham Institute, Cambridge, UK).

### RNA-sequencing bioinformatics

The raw RNASeq reads (Fastq files) for each sample were aligned to the Mus musculus 10 reference genome (https://www.ncbi.nlm.nih.gov/assembly/GCA_001269945.2) using STAR v2.4.0^60^ with default parameters, using the DNAnexus platform. After alignment, estimation of transcript abundance measures as fragments per kilobase of exon per million aligned fragments (FPKM) values was performed using Cufflinks in the Tuxedo protocol, and FPKM values were used for generation of all heatmaps. For pairwise comparisons aligned reads were then analyzed for differential gene expression utilizing the limma software package^61^ run through the BioJupies analysis package with default parameters^62^. Statistical significance was set at a threshold of FDR-correct *p* < 0.05. Significantly regulated gene lists were separately entered into G:Profiler^63^, and significantly regulated pathways were identified using an FDR corrected *p* < 0.05. Identification of predicted transcription factors was done using the Enrichr software package querying the Chea database^44–46^. Selected targets for display were those with the largest difference in transcription factor enrichment score between the H_2_O-Mor and Abx-Mor groups. Histone modifiers were identified using the Database of Epigenetic Modifiers (dbEM)^47^ and every identified target was separately confirmed to have epigenetic activity in mouse using UniProt gene database before inclusion.

### Quantitative PCR

Mice from cages receiving H_2_O, Abx, SCFA, or SCFA+Abx (n = 5 / group) were euthanized by rapid decapitation and NAc was removed as described above. Tissue was homogenized on ice and RNA was extracted from whole tissue using RNeasy kit (Qiagen) with optional DNase digestion step per assay instructions. Total RNA was quantified using NanoDrop1000 and was converted to cDNA using iScript cDNA synthesis kit (Bio-Rad) on a SimpliAmp Thermal cycler (Applied Biosystems). For quantitative PCR, samples were run in triplicate; each reaction contained 3 μg cDNA with 10 μM forward primer, 10 μM reverse primer, and 5 μl SYBR green master mix (Applied Biosystems). Reactions were run for 40 cycles on a QuantStudio5 (Applied Biosystems). For list of primer sequences, see **Supplemental Table 7**. Fold change values were calculated using the ΔΔCt method, with *Hprt1* as the loading control.

### Statistical analysis

Statistical significance for observed taxonomic units and Shannon index were analyzed using a 2 x 2 between subjects ANOVA with drug treatment and drink type as fixed factors. To determine differences in phylum abundance, individual t-tests were conducted between the control group (H_2_O-Sal) and every other group. SCFA levels in caecum were converted to fold change and significance was determined using a 3 x 2 mixed ANOVA with metabolite as a fixed within-subjects factor and drink type as a fixed between-subjects factor. Locomotor sensitization over days was analyzed using a 5 x 2 mixed ANOVA with day as a fixed within-subject factor and drink type as a fixed between-subject factor. Persistence of sensitization was analyzed using an independent samples t-test. For morphine place preference with two conditioning days, preference score was analyzed using a 3 x 2 between-subjects ANOVA with morphine dose and drink type as fixed factors. For three conditioning days, significance of preference score was calculated using an independent samples t-test. Statistical significance for qPCR was determined using separate 2 x 2 between-subjects ANOVAs for each target, with Abx (presence or absence) and SCFA (presence or absence) as fixed factors. For CPP after SCFA replenishment, data was analyzed using a 2 x 2 between subjects ANOVA with Abx (presence or absence) and SCFA (presence or absence) as fixed factors. For all relevant studies, Tukey’s HSD post hoc analysis was used to make pairwise comparisons in the presence of a significant interaction and additional planned comparisons were conducted between Abx and Sal groups as noted in the text. All t-tests are two-tailed. Unless otherwise stated, *p* values less than 0.05 were deemed statistically significant.

## Author Contributions

D.D.K. and R.S.H. designed the experiments. R.S.H., N.L.M., T.J.E., K.R.M., A.T.O. & D.D.K. performed experiments. R.S.H., T.J.E., K.R.M., A.T.O. & D.D.K. analyzed data. R.S.H. & D.D.K. wrote the manuscript. All authors provided critical edits and feedback of the finalized manuscript.

## Data Availability

All RNA-sequencing files will be uploaded to the publicly available Gene Expression Ombibus server upon publication. All data in this paper will be made available upon reasonable request.

## Acknowledgements

This work was supported by NIDA grants DA044308 & DA051551 to D.D.K. as well as by NARSAD Young Investigator Awards to R.S.H and D.D.K. Morphine sulfate was provided by the NIDA Drug Supply Program. We acknowledge the Microbial Culture & Metabolomics Core of the PennCHOP Microbiome Program for performing targeted metabolomics analyses

