## Supplemental Figures 1-2 for "The gut microbiome and its metabolites are necessary for morphine reward"

### Figure S1.

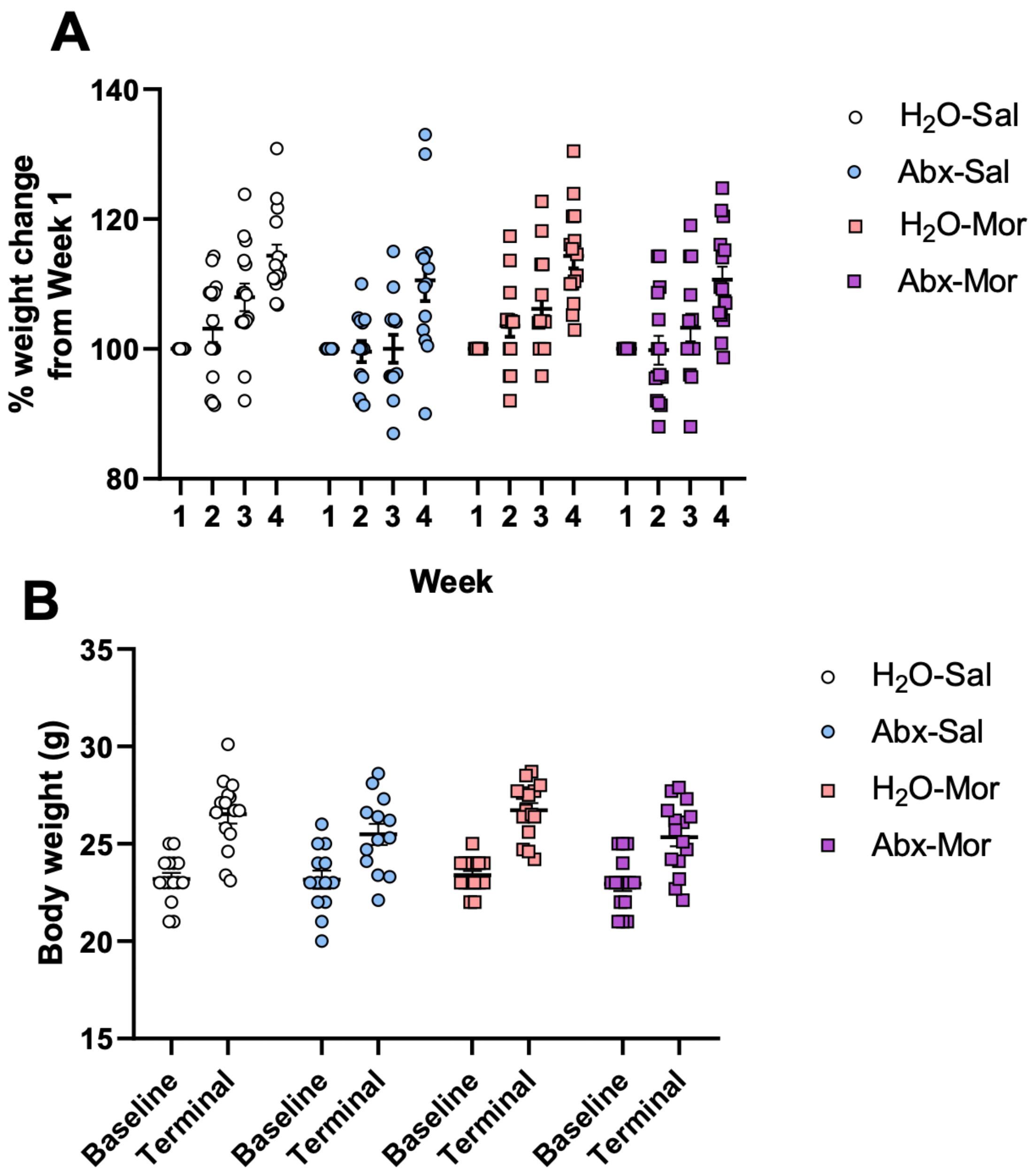


**Supplemental figure 1. Abx alone or in combination with morphine does not affect body weight.** (A) Percent weight gain in mice over 4 weeks calculated as (body weight at specified week) / (body weight immediately before Abx start) x 100. A 4 x 4 mixed ANOVA with timepoint as a within-subject fixed effect and treatment as a between-subjects fixed effect identified a main effect of time only ($F_{(3, 162)} = 60.12$, $p < 0.0001$), but no effect of treatment ($F_{(3, 54)} = 2.05$, $p = 0.12$) and no interaction ($F_{(9, 162)} = 1.09$, $p = 0.37$). (B) Body weight at the start of the experiment and at the end. A 4 x 2 mixed ANOVA with timepoint as a within-subject fixed factor and treatment as a between-subjects fixed factor found a significant effect of time ($F_{(1, 54)} = 137.1$, $p < 0.0001$), but no effect of treatment ($F_{(3, 54)} = 1.28$, $p = 0.15$) or interaction ($F_{(3, 54)} = 1.28$, $p = 0.29$). N = 13-15 / group, same mice in figure panel A and B.

### Figure S2.

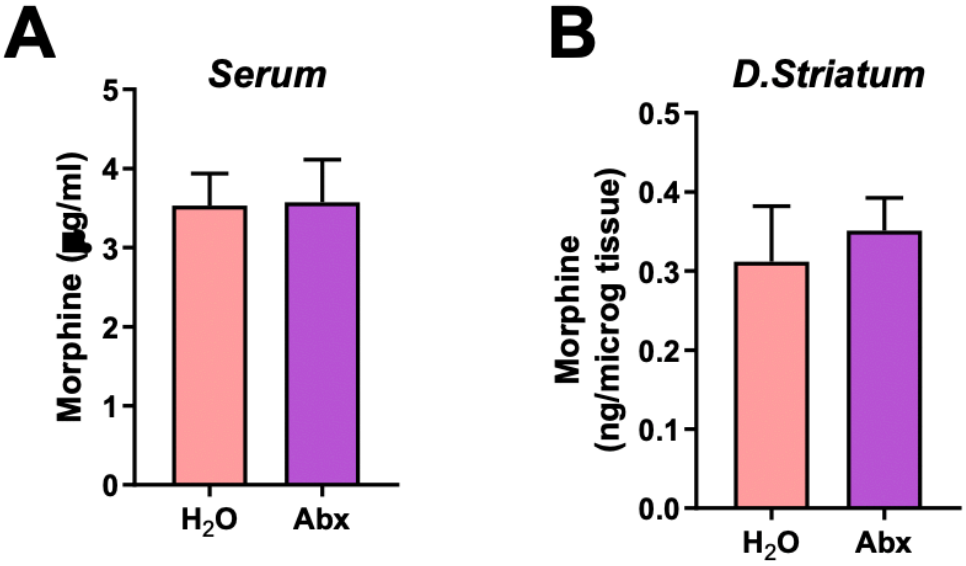


**Supplemental figure 2. Abx does not affect morphine metabolism or brain penetrance.** Levels of morphine in blood (A) and dorsal striatum (B) did not differ between $H_2O$ and Abx-treated mice. A two-tailed independent t-test serum: $t_{(8)} = 0.06$, $p = 0.95$; dorsal striatum: $t_{(8)} = 0.48$, $p = 0.65$. n = 5 / group

**Supplemental Table Legends**

**Table S1 – Bacterial Phyla** Table listing relative bacterial phyla composition derived from 16S sequencing experiments. Each grouping has individual sample columns with mean data in the right-most column. The *p* value of each sample relative to control and the fold change vs control is listed for each experimental conditoin.

**Table S2 – PICRUSt Data** Enrichment scores for individual bacterial functional molecular pathways are shown with individual values of each sample presented and statistical comparisons included in the right-most columns.

**Table S3 – RNA Seq DEGs** Tabular listing of all genes with an FDR corrected p value < 0.05 for each pairwise comparison from the nucleus accumbens RNA-sequencing dataset. Log2 fold change, Average expression, *p* value, and adjusted *p* value are shown for each pairwise comparison.

**Table S4 – Gene Ontology Analyses** Tabular formulation of gene ontology outputs produced by G:Profiler for all pairwise comparisons in the nucleus accumbens RNA-sequencing dataset.

**Table S5 – Chea Transcription Factor Enrichment** Output from the Enrichr analysis pathway predicting transcription factor binding enrichment based on the Chea dataset. Data from all mouse transcription factors is included.

**Table S6 – GO Terms from Genes Regulated in both Morphine Groups** G:Profiler output from gene list of those that were statistically significantly regulated in both the H2O-Mor and Abx-Mor groups.

**Table S7 Primers** – Primer pairs used for quantitative PCR analyses.
